# Response learning confounds assays of inhibitory control on detour tasks

**DOI:** 10.1101/838078

**Authors:** Jayden O. van Horik, Christine E. Beardsworth, Philippa R. Laker, Mark A. Whiteside, Joah R. Madden

**Affiliations:** Centre for Research in Animal Behaviour, Psychology, University of Exeter, UK

**Keywords:** Cylinder Task, Detour Task, Executive Functions, Motor Routine

## Abstract

The ability to inhibit prepotent actions towards rewards that are made inaccessible by transparent barriers has been considered to reflect capacities for inhibitory control (IC). Typically, subjects initially reach directly, and incorrectly, for the reward. With experience, subjects may inhibit this action and instead detour around barriers to access the reward. However, assays of IC are often measured across multiple trials, with the location of the reward remaining constant. Consequently, other cognitive processes, such as response learning (acquisition of a motor routine), may confound accurate assays of IC. We measured baseline IC capacities in pheasant chicks, *Phasianus colchicus*, using a transparent cylinder task. Birds were then divided into two training treatments, where they learned to access a reward placed behind a transparent barrier, but experienced differential reinforcement of a particular motor response. In the Stationary-Barrier treatment, the location of the barrier remained constant across trials. We therefore reinforced a fixed motor response, such as always go left, which birds could learn to aid their performance. Conversely, we alternated the location of the barrier across trials for birds in the Moving-Barrier treatment, and hence provided less reinforcement of their response learning. All birds then experienced a second presentation of the transparent cylinder task to assess whether differences in the training treatments influenced their subsequent capacities for IC. Birds in the Stationary-Barrier treatment showed a greater improvement in their subsequent IC performance after training compared to birds in the Moving-Barrier treatment. We therefore suggest that response learning aids IC performance on detour tasks. Consequently, non-target cognitive processes associated with different neural substrates appear to underlie performances on detour tasks, which may confound accurate assays of IC. Our findings question the construct validity of a commonly used paradigm that is widely considered to assess capacities for IC in humans and other animals.

## INTRODUCTION

Inhibitory Control (IC) is the ability to refrain prepotent responses and delay gratification (Diamond, 2013). Importantly, IC is central to the self-regulation of behaviours (Miyake & Friedman, 2012), with deficits linked to numerous pathological disorders in humans (Moffitt et al., 2011). Assays of IC, using transparent barriers, are also frequently used in studies of animal cognition (Kabadayi, Bobrowicz, & Osvath, 2018; MacLean et al., 2014). Transparent barriers are considered to evoke IC as they restrict prepotent responses towards a visible, goal placed behind the barrier (Diamond, 1981). Many subjects show initial impairments in their ability to inhibit prepotent responses, as their attempts to obtain a goal are obstructed by the barrier. With subsequent experience of the task, subjects may however improve their ability to inhibit these prepotent responses and instead detour around the barrier to obtain the goal (van Horik et al., 2018). These findings suggest that other processes of learning may mediate performances across repeated trials on these tasks, potentially confounding reliable assays of IC. Accordingly, controlled studies, using animal models, suggest that the cognitive constructs that underlie performances on some commonly used IC tasks remain unclear (van Horik et al., 2018; Völter, Tinklenberg, Call, & Seed, 2018).

A broad comparative study involving 567 individuals from 36 species found superior performances on IC tasks among anthropoid apes, leading to the notion that large absolute brain size was a good predictor of IC capacity (MacLean et al., 2014). However, subtle differences in test procedures have recently revealed that numerous species show IC performances that are comparable to those anthropoid apes reported by MacLean and colleagues (2014), even despite possessing a relatively smaller absolute brain size (corvids: Jelbert, Taylor, & Gray, 2016; Kabadayi, Taylor, von Bayern, Auguste, & Osvath, 2017; Stow, Vernouillet, & Kelly, 2018; great tits: Isaksson, Utku Urhan, & Brodin, 2018 and guppies: Lucon-Xiccato, Gatto, & Bisazza, 2017). An individual’s performance on IC tasks may also be mediated by non-cognitive processes, including differential experience with transparent barriers (van Horik et al., 2018), environmental predictability (van Horik et al., 2019) food motivation (van Horik et al., 2018), or body condition (Shaw, 2017). These findings suggest that capacities for IC, obtained from detour tasks, may suffer from task impurity. For example, individual differences in detour task performance may not be solely determined by an individual’s capacity for IC, but rather be determined by a combination of motivational and cognitive processes that confound accurate measures of IC.

Lesion studies in rodents and monkeys, alongside behavioural and neuroimaging studies in humans, reveal that orbitofrontal cortex (OFC) and lateral prefrontal cortex (lPFC) play a crucial role in regulating performances on classical IC paradigms (Diamond, 1990; Wallis, Dias, Robbins, & Roberts, 2001; but see Kabadayi et al., 2018 for review). It is likely that similar processes of IC are regulated by analogous neuroanatomical regions in birds, such as the nidopallium caudolaterale (Güntürkün, 2005). However, numerous species have been tested on different variants of detour tasks and there is little consistency in their IC performances (Brucks, Marshall-pescini, Wallis, Huber, & Range, 2017; Vernouillet, Stiles, Andrew McCausland, & Kelly, 2018a), suggesting that the construct validity of different IC tasks remains unclear (van Horik, Langley, Whiteside, Laker, Beardsworth, et al., 2018; Völter et al., 2018). It is therefore likely that performances on different detour tasks are mediated by different cognitive processes. For example, detour tasks require the inhibition of a prepotent response towards a visible reward placed behind a transparent barrier that remains in a consistent location across trials. Spatial information about the location of the reward may therefore be used to facilitate performances on detour tasks involving transparent barriers. As such, improvements in performances across trials on detour tasks may be facilitated by cognitive processes associated with the visual location of the reward, and thus involve neural substrates that are unrelated to IC *per se*. Learning the location of a reward may then be facilitated by cues in the environment, such as landmarks (i.e. *place* learning) or reinforcement of fixed motor responses, such as “turn left to access the reward” (i.e. *response* learning) (Gibson & Shettleworth, 2005; Tolman, Ritchie, & Kalish, 1946). The use of allocentric processes in spatial navigation may be determined by manipulating the location of the test apparatus or the surrounding landmark cues. Conversely, egocentric processes may be determined by presenting subjects with “Shortcut” trials, in which fixed motor responses can revealed by the perseverance of detour behaviour in the absence of the transparent barrier (Thorndike, 1911; but see Kabadayi, Bobrowicz, et al., 2018 for review). Importantly, both *place* and *response* learning are subserved by different neural substrates, the hippocampus and the striatum [caudate] respectively (Kesner, Bolland, & Dakis, 1993; Mcdonald & White, 2013; McDonald & White, 1994; Packard, Hirsh, & White, 1989; White & McDonald, 2002). Successful performances on detour tasks may therefore rely on multiple, different, cognitive processes or neural substrates, which may further confound accurate assays of IC.

In this study we attempt to clarify the role of response learning in detour task performance, and hence improve the accuracy of IC assays. Pheasant chicks, *Phasianus colchicus*, provide an excellent opportunity to investigate the processes of learning that underlie IC performance, as large numbers of birds can be hatched on the same day, reared and tested under controlled experimental conditions, and they readily engage with typical IC apparatuses (Meier et al., 2017; van Horik et al., 2018; van Horik, Langley, Whiteside, Laker, & Madden, 2018). We measured baseline levels of IC by presenting birds with a transparent cylinder task containing a food reward (MacLean et al., 2014; van Horik et al., 2018). Birds were then randomly assigned to one of two treatment groups, in which they were trained to access a food reward that was positioned behind a transparent barrier. The location of the barrier remained fixed across trials for birds in the Stationary-Barrier treatment but alternated in location across trials for birds in the Moving-Barrier treatment. All birds were then retested on the cylinder task. If response learning confounds accurate assays of IC, we expect performances between the first (baseline) and second (retest) presentations of the cylinder task to differ according to the experimental treatments each group received. Specifically, we expect birds in the Stationary-Barrier treatment to show greater improvements on subsequent IC tasks as we reinforced the acquisition of a behavioural response (motor routine), in relation to the barrier, to facilitate their performances. Conversely, we expect birds in the Moving-Barrier treatment, which adopted inconsistent behavioural responses, to show no improvement in their performances when retested on the cylinder task. To further investigate the persistence of a motor routine, we also presented all birds with a single Shortcut trial, after the Response Learning trials, in which the transparent barrier was absent. The performances of birds that unnecessarily persisted in their detour responses in the absence of the transparent barrier were considered to further reflect a fixed motor behaviour, rather than responding appropriately to the new paradigm (Verbruggen, Best, Bowditch, Stevens, & McLaren, 2014; but see Kabadayi, Bobrowicz, et al., 2018). We tested whether the use of the shortcut differed between the Moving-Barrier and Stationary-Barrier treatments, and whether birds that used the shortcut made fewer overall pecks, and hence showed greater IC, than birds that failed to respond to the shortcut. To determine whether performances on each task could be explained by non-cognitive traits that may influence a subject’s motivation to interact with an apparatus, as has been found in other studies of IC (Shaw, 2017; van Horik et al., 2018), we also assessed whether IC performances were influenced by subjects’ sex and/or body condition. We also measured their motivation to interact with the test apparatus by recording latencies to acquire a freely available mealworm (Free-Worm) that was positioned adjacent to each apparatus.

## METHODS

### Subjects and Housing

One hundred and twenty-six pheasant chicks were hatched in incubators on the same day, randomly assigned into four replicated pens, and reared from one day old between 24 May and 25 July 2018 (63 days old). All birds were identifiable from individually numbered wing tags, supplied with commercial pheasant feed (Keepers’ Choice) and water *ad libitum*. For the first 2 weeks of life birds were housed in one of four heated pens (2m × 2m) after which they had access to an adjacent covered enclosure (1m × 4m) and an outdoor run (4m × 12m).

### Procedure

Day-old chicks were habituated to human observation and shaped for the first five days of their lives, using mealworm rewards, to individually enter an experimental chamber (0.75m × 0.75m) placed adjacent to their pens. After shaping, all birds willingly entered the experimental chamber. During experimental test trials, an experimenter opened a sliding door that allowed the birds to individually enter the experimental chamber at will. After entering, the sliding door was closed, and the subject’s performance was recorded by an observer. All birds were tested individually while visually isolated from other test subjects. After testing, subjects were released into the outdoor run. Subjects that failed to engage with the tasks within five minutes from entering the experimental chamber were released and excluded from analyses. Specific protocols for each task will be described in detail below (sections 1-5; see also Figure 1). Subjects first participated in a Baseline IC Task, involving Opaque (training) and Transparent Cylinders (test). All birds in a pen were then assigned to one of two experimental treatments, in which birds were trained to acquire a reward placed behind a transparent barrier. For the Stationary-Barrier treatment group, the location of the barrier and reward remained in a fixed location across trials. Hence, we reinforced consistent behavioural responses, which they could use to facilitate their retrieval of the reward. Conversely, the location of the barriers and reward alternated between the left and right of the experimental chamber for birds in the Moving-Barrier treatment group. Hence, consistent behavioural responses were unavailable to these birds and could not be learned to facilitate their acquisition of the reward. Birds were then presented with a single Shortcut trial, to determine whether they persisted in their detour responses in the absence of the transparent barrier. Finally, all birds were retested on the Transparent Cylinder task (identical to the Baseline Cylinder task) to determine whether the different treatments experienced during training influenced their subsequent performances.

**Figure 1.**
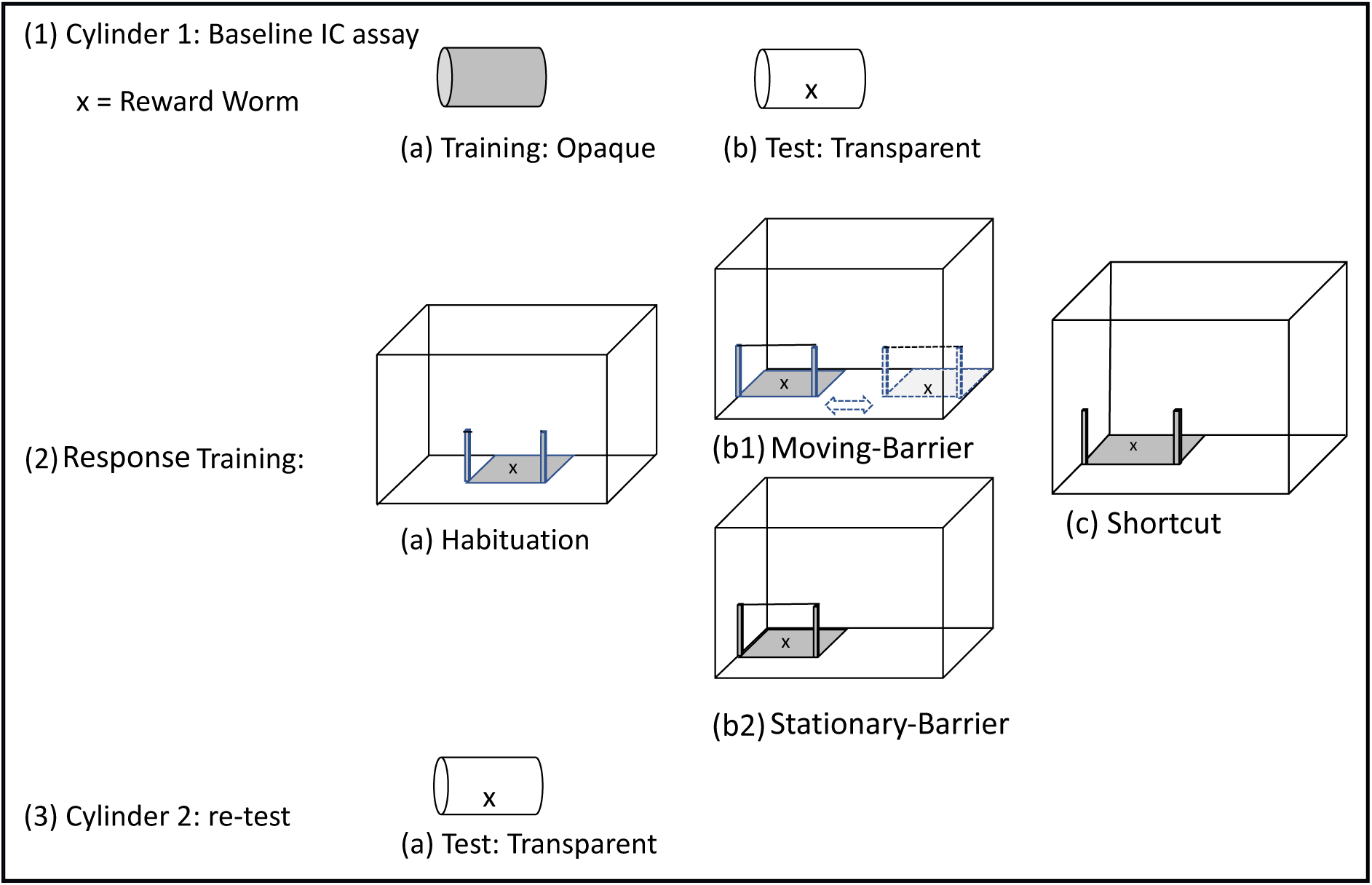
Schematic order of procedures for training and testing apparatuses. Subjects began with (1) Cylinder 1, where they participated in Baseline assays of IC using (a) training and (b) test apparatuses, and proceeded to (2) Response Training, where all birds participated in (a) Habituation trials, after which they were assigned to (b1) Moving-Barrier and (b2) Stationary-Barrier treatments and then all birds were presented with a (c) Shortcut trial. Cubes represent the experimental chamber and the relative position of each apparatus. Finally, all birds were retested on (3) Cylinder 2, (as in 1b) to determine how Response Training treatments influenced subsequent inhibitory control performance.

#### 1) Cylinder 1: Do transparent cylinders evoke prepotent responses?

We presented birds with a Cylinder detour task that is commonly used to assess capacities for inhibitory control in a variety of animals (MacLean et al., 2014). Birds first participated in five trials on an opaque training apparatus and then subsequently participated in two test trials on a transparent variant of the apparatus. On all trials, the cylinder apparatus was presented in the centre of the experimental chamber and adjacent to the subject, so the open ends were not directly in view. We positioned the Cylinder task in the centre of the testing chamber to differentiate the requirements of the Cylinder task and the subsequent Barrier task. Hence the reinforcement of the motor routine was in relation to the barrier (task specific) rather than the reinforcement of a specific route inside the testing chamber that could be adopted as a heuristic rule across tasks. The opaque training apparatus was used to habituate subjects to a novel apparatus and ensure that they could access a mealworm reward that was placed inside the cylinder before participating in the transparent test condition. Apart from transparency, and hence the visibility of the reward, the training and test apparatuses were identical. As the mealworm reward was clearly visible within the cylinder during the test condition, subjects had to inhibit their prepotent attempts to acquire the reward directly through the transparent cylinder and instead detour around to the open end of the cylinder to access the reward, as they had previously learned during the opaque training condition. However, as subjects had no experience with transparent barriers prior to testing, we acknowledge that birds would require at least one error (peck) to determine that the transparent cylinder was impenetrable. Each cylinder was 5cm diameter × 12cm long and mounted on a white 20cm × 20cm base for stability. For each trial we recorded (i) Approach latency (s) from entering the experimental chamber to consuming a freely available mealworm (hereafter Free-Worm) placed in front of the apparatus, (ii) the number of Pecks (incorrect attempts) each individual directed towards the transparent barrier before acquiring the mealworm inside the cylinder as a measure of their inhibitory control. Birds participated in two opaque training trials per day, one in the morning (0830-1230) and one in the afternoon (1400-1800), between 19-22 June 2018 (27-30 days old). To assay improvements in IC performances across trials, we presented all birds with two transparent test trials, one in the afternoon on 22 June 2018 (30 days old) and one in the morning on 25 June 2018 (33 days old).

#### 2) Habituation and Response Training: moving vs stationary transparent barriers

After completing the Baseline IC Assay, but immediately prior to Response Training, all birds received four habituation trials in which they encountered the Response Training apparatus without a transparent barrier. During these habituation trials the frame of the apparatus was placed in the centre of the experimental chamber and was comprised of a wooden base (40cm long × 25cm wide), with a wooden post (30cm high) at either end, between which the transparent barrier (40cm wide × 30cm high) would be subsequently attached during Response Training trials. For each trial we placed 5 mealworms inside a white lid (5cm diameter) with a 1cm lip so that the worms could be seen but not escape. During habituation trials the lid was positioned in the centre of the apparatus. Subjects therefore had to step onto the wooden base to acquire the reward. The purpose of the habituation trails was to reduce any neophobic responses towards the apparatus and to reinforce birds to approach the reward between the two wooden posts. For each trial, we recorded each subject’s latency from entering the experimental chamber to consuming the first mealworm inside the white lid. Birds participated in three habituation trials on 25 June 2018 (33 days old) and one habituation trial in the morning on 26 June 2018 (34 days old).

After completing the habituation trials, a transparent barrier was fixed to the wooden posts and prevented birds from approaching the reward directly. Birds were randomly assigned to one of two Response Training treatments in which they could access rewards that were placed behind the transparent barrier (Figure 1). Birds experienced 10 Response Training trials in which the location of the barrier either moved or was stationary depending on the treatment. Birds participated in two Response Training trials on 26 June 2018 (34 days old), and four trials per day on 27 and 28 June 2018 (35 and 36 days old). In the Stationary-Barrier treatment, the barrier was consistently located either on the left or right of the experimental chamber (counterbalanced across individuals). A fixed behavioural response was therefore consistently reinforced for birds in the Stationary-Barrier treatment. In the Moving-Barrier treatment, the barrier location alternated between the left- and right-hand side of the experimental chamber across trials. Inconsistent (as opposed to consistent) behavioural responses were therefore reinforced for birds in the Moving-Barrier treatment. For each treatment, the lid containing the mealworms was consistently positioned at the far end of the apparatus (see Figure 1). To access the reward, subjects had to inhibit directly approaching the reward and instead detour around the barrier which could only be accessed from one side. During Habituation and Training trials, we recorded the subject’s latency from entering the chamber to acquiring a Reward-Worm placed behind the barrier arms. During Training trials, we recorded the number of incorrect attempts (Pecks) to acquire the Reward-Worm through the transparent barrier.

#### 3) Shortcuts: Do birds persist in their detour behaviours in the absence of the barrier?

After completing the 10 Response Training trials, all birds were presented with a single “Shortcut” trial on 28 June 2018 (36 days old) that was identical to the initial habituation trial, where the transparent barrier was absent. The Shortcut apparatus was positioned in the same or alternating location as in the Response Training trials for each respective treatment group. During Shortcut trials, we recorded the subject’s latency from entering the chamber to acquiring a Reward-Worm placed behind the barrier arms.

#### 4) Cylinder 2: Do non-target cognitive processes influence IC performance?

After completing the Shortcut trial, all birds were retested with the transparent Cylinder task (using identical procedures as in the IC Baseline assay), to determine whether Response Training influenced their subsequent capacities for IC. Birds experienced one trial on this task between 09:30-12:30 hrs on 29 June 2018 (37 days old).

#### 5) Do non-cognitive/motivational processes influence task performances?

To determine whether IC performances were influenced by non-cognitive factors, we positioned a freely available mealworm (Free-Worm) adjacent to each test apparatus. The purpose of the Free-Worm was (i) to standardise the approach direction of each subject, (ii) to ensure subjects were motivated by food rewards and (iii) determine whether approach latencies differed across trials, which may suggest performances were influenced by neophobic responses towards an apparatus. On 20 July 2018, after birds had participated in all tests, we recorded each individuals’ mass (Slater Super Samsom spring balance – precision 5 g), and tarsus length (callipers – precision 0.1 mm), to determine their body condition (mass/tarsus^3^). Birds in poor body condition (low scores) were considered to be more food-motivated than birds in good body condition (high scores). As male pheasants are larger than female pheasants (Whiteside, van Horik, Langley, Beardsworth, & Madden, 2018), differences in growth rates may lead to motivational differences, and we have previously found these to differentially influence participation on cognitive tests (van Horik, Langley, Whiteside, & Madden, 2017). We therefore used plumage features to visually identify the sex of each individual at 10 weeks old.

### Inclusion/exclusion of subjects for analyses

To ensure that experience on each task was standardised across subjects, we only included birds that participated in and acquired the Reward-Worm on all trials for all tasks. Hence, all birds included in this study experienced: five opaque cylinder training trials; two transparent cylinder test trials; four no-barrier habituation trials; 10 Response Training trials; one Shortcut trial; and one transparent cylinder retest trial. Sixty-two subjects met all these criteria (Moving-Barrier: 16 males; 9 females; Stationary-Barrier: 20 males; 17 females). Birds that were excluded either pecked at the apparatus but failed to acquire the mealworm reward, or failed to interact with the apparatus. Birds in the former category were excluded because we could not ensure equal competency in retrieving the reward. Hence, a failure to retrieve the reward may be due to inexperience rather than poor IC. Birds in the latter category were excluded because we could not obtain accurate assays of performance, which were likely due to neophobic responses towards the apparatus.

### Statistical analysis

We used Generalised Linear Mixed Models (GLMMs), using the lme4 package (Bates, Maechler, Bolker, & Walker, 2015) in R (R Development Core Team, 2014) to assess performances on all tasks, excluding the Shortcut trial and improvements between the Cylinder 1 and Cylinder 2 tasks, were we used Generalised Linear Models (GLM). To determine whether the transparency of the cylinder evoked prepotent responses, we compared the number of Pecks (errors) that subjects made when attempting to acquire the mealworm (Reward-Worm) between the Opaque and Transparent Cylinder tasks. We assessed learning on the transparent Cylinder task by comparing pecks across trials. Latencies from entering the experimental chamber to acquiring a Reward-Worm that was positioned inside each apparatus were used as performance measures during the No-Barrier Habituation trials because there was no barrier to peck at. Latencies to acquire the Reward-Worm, as well as Pecks to the transparent barriers were used as performance measures during Response Training. We used a Binomial Test (set at 0.5) in SPSS (IBM Corp, 2013) to determine whether birds persisted in their detour behaviours by avoiding an absent barrier during Shortcut trials, or whether they used the Shortcut and went through the barrier arms to access the mealworm reward. To determine whether the Response Training treatments had differential influences on subsequent IC performances, we subtracted the number of Pecks that each individual made on their second trial of the Baseline Transparent Cylinder task (Cylinder 1) from the number of Pecks they made when retested on the Transparent Cylinder task after Response training (Cylinder 2). Hence, a negative score indicates a reduction in Pecks (errors) when retested and we considered this to indicate improvement in performance. We also assessed whether performances on the Shortcut trials predicted improvements in pecks between the Cylinder 1 and Cylinder 2 tasks. Pecks were assessed using a poisson error distribution and Reward-Worm latencies were assessed using a gaussian error distribution (lmer). Depending on the task (see Table 1), we assessed whether our performance measures were influenced by the following predictor variables: Free-Worm latency, Sex (female = 0; male = 1), Body Condition, Treatment (Moving-Barrier = 1 vs Stationary-Barrier = 0) and Trial Number, Shortcut (around barrier = 0; through barrier = 1). When using GLMMs, we included bird as a random effect to control for pseudoreplication, and included an observational-level random effect to control for overdispersion (Harrison, 2014).

**Table 1.** Predictor variables and model outputs for GLMMs (Pecks: models 1, 2, 3b and Reward-Worm latencies: model 3a,c), and GLM (Reward-Worm latencies: model 4; Pecks: model 5). Estimates ± SEM are presented with their corresponding Chi Squared (X^2^) and significance values (p). n/a = variable not included in analysis.

| Models | Free-Worm | Sex | Body Condition | Treatment | Trial |
| --- | --- | --- | --- | --- | --- |
| 1) Cylinder 1: Opaque vs Transparent | 0.018 ± 0.038;<br>X<sup>2</sup> = 0.222, p = 0.638 | 0.030 ± 0.262;<br>X<sup>2</sup> = 0.013, p = 0.301 | 0.206 ± 5.780;<br>X<sup>2</sup> = 0.305, p = 0.581 | n/a | -0.231 ± 1.366;<br>X<sup>2</sup> = 172.798, p < 0.001 |
| 2) Cylinder 1: Transparent<br>[Improvement across trials] | 0.022 ± 0.028;<br>X<sup>2</sup> = 0.605, p = 0.437 | 0.304 ± 0.163;<br>X<sup>2</sup> = 3.43, p = 0.064 | 2.376 ± 3.638;<br>X<sup>2</sup> = 0.424, p = 0.515 | n/a | -0.560 ± 0.156;<br>X<sup>2</sup> = 12.167, p < 0.001 |
| 3a) No-Barrier Habituation | n/a | -2.638 ± 4.531;<br>X<sup>2</sup> = 0.355, p = 0.551 | -53.045 ± 101.676;<br>X<sup>2</sup> = 0.285, p = 0.593 | -0.961 ± 4.491;<br>X<sup>2</sup> = 0.049, p = 0.825 | -11.993 ± 1.459;<br>X<sup>2</sup> = 57.935, p < 0.001 |
| 3b) Response Training: Pecks | n/a | -0.144 ± 0.207;<br>X<sup>2</sup> = 0.48, p = 0.487 | -8.209 ± 4.678;<br>X<sup>2</sup> = 2.98, p = 0.084 | 1.146 ± 0.200;<br>X<sup>2</sup> = 363.73, p < 0.001 | -0.600 ± 0.029;<br>X<sup>2</sup> = 363.73, p < 0.001 |
| 3c) Response Training: Reward-Worm | n/a | -9.426 ± 4.890;<br>X<sup>2</sup> = 3.85, p = 0.050 | -21.760 ± 109.206;<br>X<sup>2</sup> = 0.042, p = 0.837 | 12.746 ± 4.784;<br>X<sup>2</sup> = 7.159, p = 0.007 | -8.183 ± 0.588;<br>X<sup>2</sup> = 166.764, p < 0.001 |
| 4) Shortcut<br>[Treatment = through vs around barrier] | n/a | -2.660 ± 5.098;<br>X<sup>2</sup> = -0.522, p = 0.604 | -48.186 ± 116.806;<br>X<sup>2</sup> = -0.413, p = 0.681 | 7.094 ± 6.140;<br>X<sup>2</sup> = 1.155, p = 0.253 | n/a |
| 5) Cylinder 2: Retest<br>[Post training improvement] | 0.067 ± 0.569;<br>X<sup>2</sup> = 0.015, p = 0.902 | -4.368 ± 4.647;<br>X<sup>2</sup> = 0.954, p = 0.329 | -24.668 ± 103.093;<br>X<sup>2</sup> = 0.062, p = 0.803 | 18.496 ± 4.533;<br>X<sup>2</sup> = 15.886, p < 0.001 | n/a |

### Ethics

All work was approved and conducted under Home Office licence PPL 30/3204 and approved by the University of Exeter Animal Welfare Ethical Review Board.

## RESULTS

### 1) Cylinder 1: Do transparent cylinders evoke prepotent responses?

Only two of 62 birds in this study made no errors on their first trial of the transparent Cylinder task, and all birds pecked at least once at the transparent cylinder on their second trial. Hence, we consider that all birds had experience that the transparent cylinder was impenetrable. Birds pecked more frequently, and hence made more incorrect attempts to acquire the mealworm placed inside the cylinder, when the apparatus was transparent rather than opaque (Table 1, model 1: Opaque Cylinder trial 5 mean pecks = 0.629 ± 0.282 SEM; Transparent Cylinder Trial 1 mean pecks = 31.161 ± 2.586 SEM).

### 2) Cylinder 1: Do baseline inhibitory control performances improve across trials?

Birds improved their Baseline IC performances across trials on the transparent cylinder task, making approximately 26% fewer pecks on their second trial compared to their first trial (Table 1: model 2).

### 6) Habituation and Response Training: moving vs stationary transparent barriers

Birds showed an improvement in their Reward-Worm latencies across the habituation trials when the transparent barrier was absent (Trial 1 mean latency 39.950 ± 6.104 SEM; Trial 2 mean latency 13.9661 ± 3.171 SEM; Trial 3 mean latency 5.212 ± 0.888 SEM; Trial 4 mean latency 2.890 ± 0.461 SEM), suggesting a reduction in neophobia towards the apparatus (Table 1: model 3a). During Response Training, birds in the Moving-Barrier treatment pecked at the transparent barrier more frequently, and took longer to acquire the Reward-Worm, than birds in the Stationary-Barrier treatment (Table 1: model 3b,c; Figure 2). Pecks and Reward-Worm latencies also decreased across trials for both treatment groups (Table 1: model 3b,c; Figure 2). Reward-Worm latencies and Pecks were unrelated to Body Condition (Table 1: model 3b,c).

**Figure 2.**
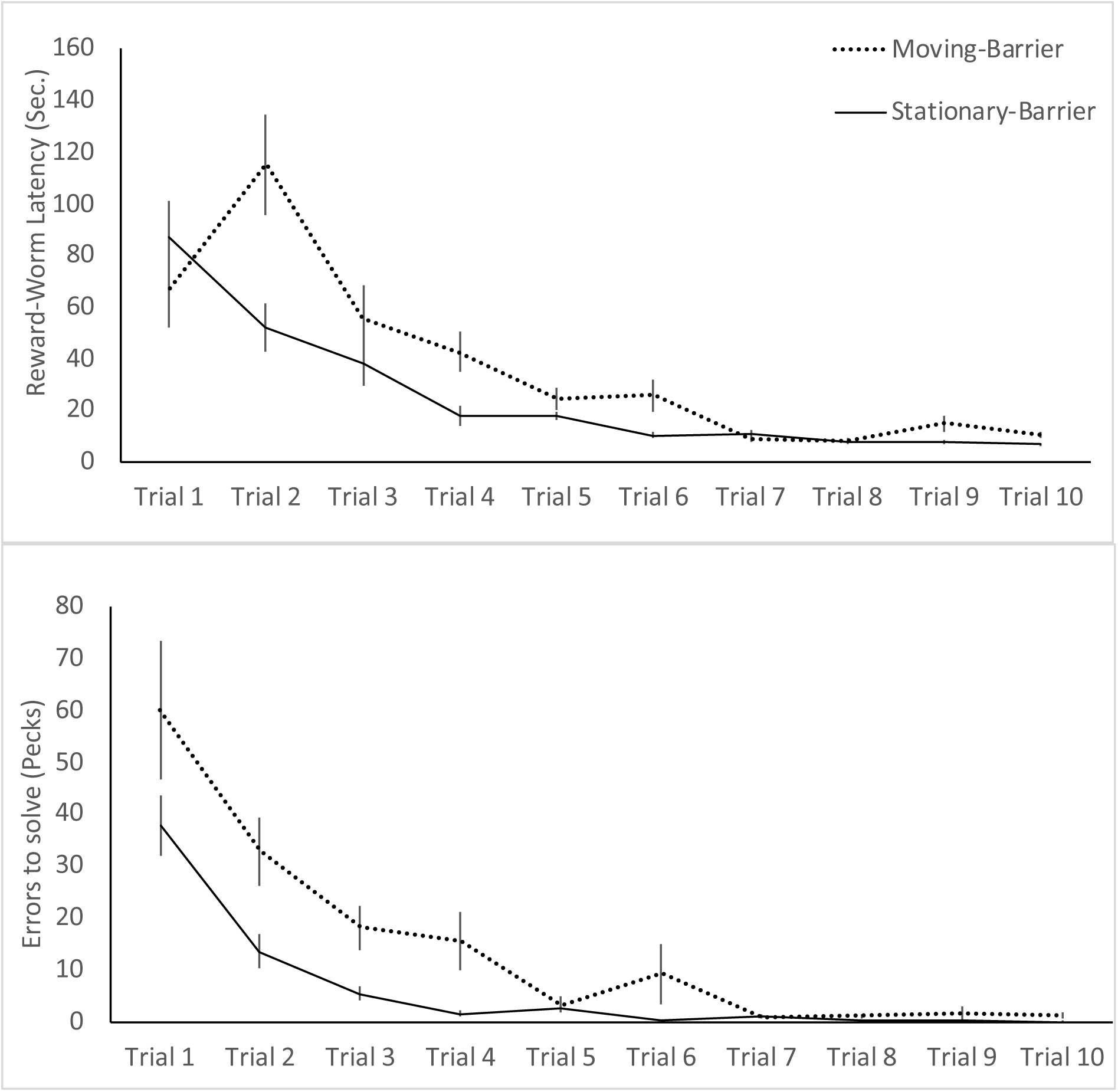
Response Training. Latencies to acquire the Reward-Worm (top) positioned behind a transparent barrier and pecks, indicating prepotent errors (bottom) across 10 trials, for birds in the Moving-Barrier (dashed line) and Barrier-Stationary (solid line) treatment groups (means ± SEM).

### 7) Shortcuts: Do birds persist in their detour behaviours in the absence of the barrier?

When the barrier was absent, birds in both treatments were more likely to go through the “Shortcut” (i.e. between the barrier arms) than detour around the absent barrier. Barrier Stationary Treatment: 26 of 37 birds (70%) went through the barrier; Binomial Test with a probability set at 0.5, p = .010. Barrier Movement Treatment: 23 of 25 birds (92%) went through the barrier; Binomial Test with a probability set at 0.5, p < .001. Improvement in errors (pecks) on the Cylinder task re-test were unrelated to whether or not birds avoided the absent barrier on the Shortcut trial (Table 1: model 4).

### 8) Cylinder 2: Do non-target cognitive processes influence IC performance?

Birds from the Stationary-Barrier treatment made approximately 58% fewer pecks when retested on the Transparent Cylinder task (after Response Training), whereas birds Moving-Barrier treatment made approximately 4% more pecks. Hence, birds from the Stationary-Barrier treatment showed a greater improvement in their IC performances (reduction in pecks relative to their baseline performance) compared to birds from the Moving-Barrier treatment (Table 1: model 5).

### 9) Do non-cognitive/motivational processes influence task performances?

Differences in performances on all tasks were generally unrelated to Free-Worm latencies, Sex or Body Condition (Table 1). However, Sex predicted Reward-Worm latencies during Response Training, with females initially taking longer to acquire the Reward-Worm than males, but with both sexes showing comparable performances after 10 Response Training trials (Table 1: model 3c; Figure 3).

**Figure 3.**
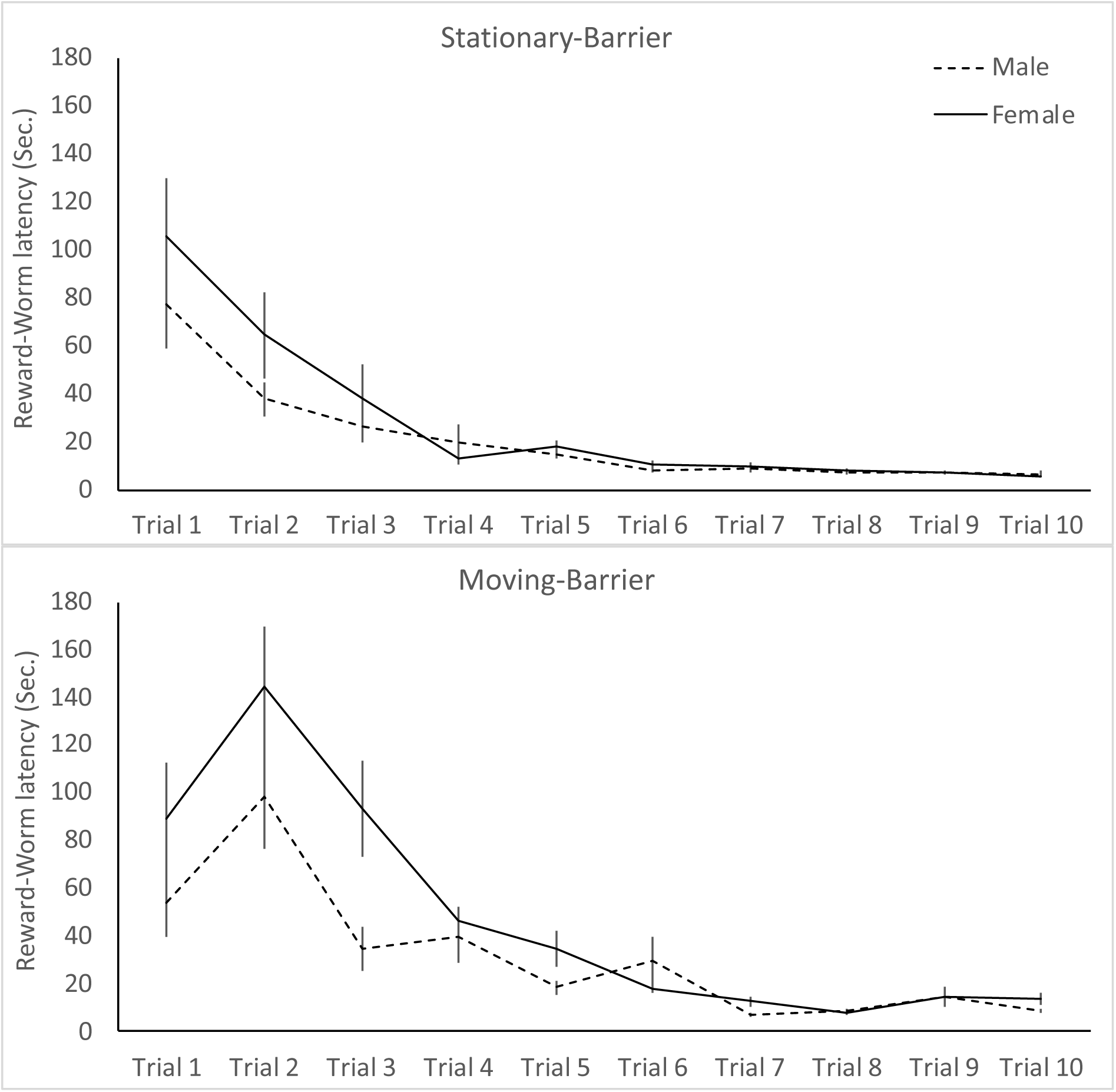
Response Training latencies (mean ± SEM) to acquire a Reward-Worm positioned behind a transparent barrier across 10 trials, for males (dashed line) and females (solid line).

## DISCUSSION

We altered inhibitory control (IC) performances of young pheasants on a transparent cylinder task, by experimentally manipulating the reinforcement of a fixed behavioural response during training on a transparent barrier task. We found that the reinforcement of a fixed behavioural response (acquisition of a motor routine) improved subsequent IC performance. These findings suggest that response learning plays an important role in facilitating successful performances on detour tasks involving transparent obstacles. Consequently, accurate assays of IC obtained from detour tasks using transparent barriers may be confounded by multiple cognitive processes that are unrelated to IC.

Capacities for IC have been considered to underlie performances on detour tasks (Diamond, 1981; Kabadayi, Bobrowicz, et al., 2017). To some extent our findings support these claims. Pheasant chicks successfully learned to extract a mealworm reward from inside an opaque cylinder, but pecked more frequently, making more incorrect attempts to acquire the mealworm, when presented with a transparent version of the apparatus. Consequently, the visibility of the mealworm inside the transparent cylinder evoked prepotent responses, which must be inhibited to acquire the reward (see Vernouillet, Stiles, Andrew McCausland, & Kelly, 2018). However, baseline IC performances on the transparent cylinder task also improved across trials, with birds making fewer erroneous pecks to acquire the mealworm reward on their second trial than compared to their first trial, as has been found in numerous other studies (Lucon-Xiccato et al., 2017; van Horik, Langley, Whiteside, Laker, Beardsworth, et al., 2018; Vernouillet et al., 2018). Moreover, latencies to acquire the mealworm reward, and pecks, also decreased across trials during response training when the reward was placed behind a transparent barrier. Although we observed an initial neophobic response towards the response training apparatus during habituation (i.e. latencies to acquire the reward decreased across trials), we consider it unlikely that improvements in IC performance across trials were due to a reduction in neophobia, as latencies to approach the apparatus did not influence IC performances. However, as birds had no prior experience with transparent barriers, an alternate explanation that could account for a decrease in errors and latencies across trials is that the number of pecks on Trial 1 was confounded by a lack of experience. Consequently, birds may have pecked more frequently on Trial 1 to explore the properties of the impenetrable transparent barrier. While this explanation is difficult to refute, all but two birds pecked at least once at the transparent apparatus during their first trial on the baseline IC task. It therefore remains possible that the physical properties of the barrier were experienced by most birds after their first peck, and that any subsequent pecks were mediated by other processes of learning and inhibitory control. Importantly, when retested on the transparent cylinder task after response training, we found a greater improvement in baseline IC performances for birds that received stronger reinforcement of a fixed behavioural response during response training (Stationary-Barrier treatment) than compared to birds that received no consistent reinforcement for behavioural responses during training (Moving-Barrier treatment). We therefore consider that improvements in performance across trials were mediated by processes of learning. Specifically, we suggest that these processes of learning were facilitated by the acquisition of a fixed motor routine, i.e. response learning (Tolman et al., 1946). However, we found that birds were more likely to use the Shortcut when the transparent barrier was absent than persist in their redundant detour behaviours. Moreover, improvements in performances on the cylinder re-test did not differ between birds that either used the shortcut or failed to respond to the shortcut.

Pecks at the transparent barrier were always directed towards the mealworm, and birds from both treatments pecked more frequently on the first trial of the barrier task than compared their preceding trials on the cylinder task. We have previously reported similar findings, in the same system, suggesting that barrier tasks may be more difficult to solve than the cylinder task (van Horik, Langley, Whiteside, Laker, Beardsworth, et al., 2018). However, van Horik and colleagues (2018) also show improvements in subsequent task performances when presented with both tasks in a counterbalanced order. These findings suggest that birds show some functional generalisation of learned affordances between barrier and cylinder tasks. Performances on the response training trials did however differ between the two treatment groups. Birds in the Stationary-Barrier treatment made fewer pecks and acquired the reward faster than birds in the Moving-Barrier treatment. While the consistent location of the barrier and reward appeared to facilitate improvements in performances of birds in the Stationary-Barrier treatment, it is possible that a violation of expectancy of the reward location contributed to increased latencies to solve the task. Interestingly, birds in the Moving-Barrier treatment also pecked more frequently at the apparatus compared to those in the Stationary-Barrier treatment. This difference in pecks between the two treatment groups was particularly evident on the first trial of the response training task, in which we might expect performances not to differ between the two treatment groups. It therefore remains possible that, by chance, birds we had randomly assigned to the Moving-Barrier treatment simply pecked more frequently than birds in the Stationary-Barrier treatment even before they had an opportunity to learn the task affordances. To test the role of motor-learning on IC performance further, subsequent studies could test whether fixed motor reinforcement facilitated particular side preferences on the cylinder task. Subsequent studies could also introduce an additional control group, where subjects receive no response training trails (of either a Moving or Stationary-Barrier). If performances were not facilitated by motor rule learning, then we might expect birds in the control group, that receive no response training, to show equivalent improvements in performances on the cylinder task re-test to those in the Stationary-Barrier treatment.

Previous studies have shown that a variety of additional factors, such as body condition (Shaw, 2017), motivation (van Horik et al., 2018), temperament (Bray, MacLean, & Hare, 2015), age (Bray, MacLean, & Hare, 2014), experience (Barrera, Alterisio, Scandurra, Bentosela, & D’Aniello, 2018; van Horik et al., 2019; van Horik, Langley, Whiteside, Laker, Beardsworth, et al., 2018; but see Fagnani, Barrera, Carballo, & Bentosela, 2016), but not neophobia (Stow, Vernouillet, & Kelly, 2018b), can influence IC performance on cylinder tasks. Age and experience could not explain the performances of pheasant chicks in the current study, as all birds were hatched on the same day and experienced the identical rearing conditions (with the exception of the response training treatments). Moreover, we found that performances on the cylinder and response training tasks were generally unrelated to our motivational (non-cognitive) measures, including latencies to acquire a freely available mealworm placed adjacent to each apparatus, body condition or sex. Relationships between body condition and performance measures should however be treated cautiously, as body condition was measured immediately prior to release and not during testing. Hence, it remains unclear whether these measures were representative during testing. We also found that females took longer than males to acquire the mealworm reward during the initial response training trials. While these differences were more pronounced among females in the Moving-Barrier treatment, differences between sexes rapidly diminished across trials. We consider it unlikely that males were less neophobic towards the response training apparatus than females, as we found no effect of sex during habituation trials, or indeed for latencies to approach any other task. Hence, these sex differences remain difficult to interpret.

Our findings align with recent studies that question the construct validity of assays of IC obtained from detour tasks (Brucks, Marshall-pescini, Wallis, Huber, & Range, 2017; van Horik et al., 2018; Vernouillet, Stiles, Andrew McCausland, & Kelly, 2018; Völter et al., 2018). Importantly, we show that performances on detour tasks administered over multiple trials may be influenced by cognitive processes unrelated to IC (Kabadayi et al., 2018; van Horik, Langley, Whiteside, Laker, Beardsworth, et al., 2018). Consequently, performances on detour tasks that are administered across multiple trials may provide inaccurate assays of IC. While it remains difficult to determine whether our experimental treatments evoked response learning, rather than some other cognitive or behavioural processes that may result from the movement of barriers, we highlight the importance of considering the influence of multiple cognitive processes when inferring capacities for IC from performances on detour tasks. To overcome these issues, we suggest future studies first establish which IC tasks reveal repeatable individual differences in performances (i.e. Cauchoix et al., 2018). We also suggest that assays of IC performance on detour tasks are obtained from a minimal number of trials to avoid multiple processes of learning. However, we acknowledge that some prior experience of transparency is necessary to provide information about the impenetrability of the barrier. We also highlight the importance of assaying personality traits (i.e. exploration) that may confound assays of performance. Future studies could further test response learning by comparing the direction that birds access the transparent cylinder before and after response training and adopt different spatial manipulations, such as altering landmark cues and the position of the test apparatus, while maintaining similar treatments as in the current study. We argue that further clarity about the neural mechanisms that underlie performances on different detour tasks is needed. Understanding these neural mechanisms will help reveal whether transparent detour tasks, that are now commonly used when testing non-human animals, can provide accurate assays of inhibitory control.

## Author Contributions

JOvH conceived and designed the experiment in discussion with JRM; JOvH, CEB, PRL, MAW collected the data; JOvH analysed data and wrote the manuscript; CEB, PRL, MAW, JRM provided comments on the manuscript.

## Data Accessibility

All data are available on Dryad

## Acknowledgements

Rothamsted Research, North Wyke hosted the rearing and release of the pheasants. Kandace Griffin and Anna Morris helped with data collection and animal husbandry.

